# A benchmarking of human mitochondrial DNA haplogroup classifiers from whole-genome and whole-exome sequence data

**DOI:** 10.1101/2021.02.11.430775

**Authors:** Víctor García-Olivares, Adrián Muñoz-Barrera, José Miguel Lorenzo-Salazar, Carlos Zaragoza-Trello, Luis A. Rubio-Rodríguez, Ana Díaz-de Usera, David Jáspez, Antonio Iñigo Campos, Rafaela González-Montelongo, Carlos Flores

## Abstract

The mitochondrial genome (mtDNA) is of interest for a range of fields including evolutionary, forensic, and medical genetics. Human mitogenomes can be classified into evolutionary related haplogroups that provide ancestral information and pedigree relationships. Because of this and the advent of high-throughput sequencing (HTS) technology, there is a diversity of bioinformatic tools for haplogroup classification. We present a benchmarking of the 11 most salient tools for human mtDNA classification using empirical whole-genome (WGS) and whole-exome (WES) short-read sequencing data from 36 unrelated donors. Besides, because of its relevance, we also assess the best performing tool in third-generation long noisy read WGS data obtained with nanopore technology for a subset of the donors. We found that, for short-read WGS, most of the tools exhibit high accuracy for haplogroup classification irrespective of the input file used for the analysis. However, for short-read WES, Haplocheck and MixEmt were the most accurate tools. Based on the performance shown for WGS and WES, and the accompanying qualitative assessment, Haplocheck stands out as the most complete tool. For third-generation HTS data, we also showed that Haplocheck was able to accurately retrieve mtDNA haplogroups for all samples assessed, although only after following assembly-based approaches (either based on a referenced-based assembly or a hybrid *de novo* assembly). Taken together, our results provide guidance for researchers to select the most suitable tool to conduct the mtDNA analyses from HTS data.

## 1. Introduction

The human mtDNA is a circular double-stranded genome of 16,569 base pairs (bp) in the hg38 reference sequence, encoding 37 genes for 13 proteins, 22 tRNAs, and 2 rRNAs. Besides its key role in diverse human diseases (DeBalsi et al., 2017; Pyle et al., 2016; West & Shadel, 2017), distinctive features such as the matrilineal inheritance, the lack of recombination, and a higher mutation rate than the nuclear genome, makes mtDNA analysis a powerful tool also for population genetics (Brotherton et al., 2013; Llamas et al., 2016; Posth et al., 2016) and forensic studies (Børsting & Morling, 2015; Just et al., 2009).

Worldwide human mtDNA diversity has been reconstructed through a genealogy of distinctive lineages, representing mtDNA sequences clustered into evolutionarily related haplotypes (a.k.a. haplogroups). They are linked to human evolutionary history and allow tracing origins and differentiation patterns across populations and over time periods (Balter, 2011; Fu et al., 2013; Gonder et al., 2007; Hajdinjak et al., 2018). As such, many haplogroups are associated with specific biogeographical ancestries (Chan et al., 2019; De Angelis et al., 2018; Maca-Meyer et al., 2001). MtDNA haplogroup classification has become an essential step to recovery ancestry and genealogical information from analyzed samples (Weissensteiner et al., 2016). One of the most popular and continuously updated repositories of mtDNA lineage relationships and nomenclature is Phylotree (http://www.phylotree.org) (van Oven & Kayser, 2009). The latest version of Phylotree (Build 17) contains more than 5,400 haplogroups and it constitutes the central reference for many bioinformatic tools to classify human mtDNA sequences.

The advent of high-throughput sequencing (HTS) technology has allowed the development of a wide range of applications, including whole-genome sequencing (WGS) and whole-exome sequencing (WES). MtDNA studies have also leveraged the power offered by HTS (Churchill et al., 2018; Schönberg et al., 2011; Vasta et al., 2009). In fact, mtDNA can be fully captured by commercial WES solutions, possibly due to the high copy number of mtDNA (Picardi & Pesole, 2012). Hence, mtDNA information can be recovered effectively from WES studies at no extra cost. Furthermore, the sequencing depth associated with WES usually allows reconstructing the entire mitogenome with high quality and detecting heteroplasmic sites (Sosa et al., 2012). Third-generation HTS, such as those based on nanopore technology (ONT, Oxford Nanopore Technologies, Oxford, UK), allows sequencing with long reads, providing an opportunity to generate full-length mtDNA sequences in single reads.

Because of the importance of recovering ancestral information and pedigree relationships of study samples, the number of bioinformatics tools developed for mtDNA haplogroup classification has increased notably in the last decade to adapt to the HTS technologies (Calabrese et al., 2014; Fan & Yao, 2011; Ishiya & Ueda, 2017; Kim et al., 2020; Navarro-Gomez et al., 2015; Röck et al., 2013; Smieszek et al., 2018; Vohr et al., 2017; Weissensteiner et al., 2016, 2020). An unmet need is a benchmarking study of their capabilities, requirements, and usability to correctly classify the mtDNA haplogroups from different source files. Based on short-read WES and WGS data from the same donors, as well as on WGS obtained from long noisy reads from a subset of them, here we present an empirical evaluation of the 11 most salient bioinformatic tools available for human mtDNA classification from HTS data.

## 2. Materials and methods

### Samples, library preparation and sequencing

The study was approved by the Research Ethics Committee of the Hospital Universitario Nuestra Señora de Candelaria and performed according to The Code of Ethics of the World Medical Association (Declaration of Helsinki).

Thirty-six DNA samples from unrelated donors of European descent were used for the study after informed consent (see supplementary **Table S1**). DNA was extracted from peripheral blood using a commercial column-based solution (GE Healthcare, Chicago, IL). DNA quantifications were performed in a Qubit dsDNA HS Assay (Thermo Fisher, Waltham, MA). All samples were sequenced in parallel using short-read WGS and WES. Library constructions were performed with Illumina preparation kits following the manufacturer’s recommendations (Illumina Inc., San Diego, CA). The Nextera DNA Prep kit was used for WGS, except for six samples that were processed by Illumina DNA Prep. The same samples were processed with Nextera DNA Exome with a 350 bp insert size as described elsewhere (Díaz-de Usera et al., 2020). The library quality control was carried out in a TapeStation 4200 (Agilent Technologies, Santa Clara, CA). Sequencing was conducted on a HiSeq 4000 Sequencing System (Illumina Inc.) at the Instituto Tecnológico y de Energías Renovables (Santa Cruz de Tenerife, Spain).

Eight of these samples had WGS based on noisy long-read data obtained at KeyGene (Wageningen, The Netherlands). Briefly, sequencing was performed on a PromethION system (ONT) for 64 h using one FLO_PR002 (R9.4.1 pore) flow cell per sample following the manufacturer’s recommendations. Basecalling was performed on the PromethION computing module using MinKNOW v1.14.2 with Guppy v2.2.2 and data preprocessing was carried out with PycoQC (Leger & Leonardi, 2019).

### Sequence processing and variant calling

Processing of short-read WGS and WES data was carried out using an in-house pipeline (see supplementary **Figure S1**) based on GATK v4.1 for WGS and GATK v3.8 for WES (McKenna et al., 2010). Reads were aligned to the GRCh37/hg19 reference genome and the mtDNA reads realigned to the revised Cambridge Reference Sequence (rCRS), GenBank NC_012920 (Anderson et al., 1981; Andrews et al., 1999), following GATK best practices for this circular genome (see supplementary **Figure S1**). This required two alignment steps: one against the canonical mitogenome reference and another against the same reference but shifted by 8,000 nucleotide positions. This generates a displacement of the mtDNA D-loop to the opposite side of the contig, allowing to better capture variants of this highly variable region. The alignments were then processed for duplicate marking and base quality score recalibration (DePristo et al., 2011). The variant calls were obtained with the “mitochondria mode” of Mutect2 GATK v4.1. This specific mode provides a better sensitivity for this genome as it shows a robust detection of very low fractions of variants once the nuclear mtDNA segments (NuMT) are filtered out. The mtDNA variants were then filtered by using FilterMutectCalls and VariantFiltration tools to keep the variants classified as PASS.

For ONT data, we first extracted all reads aligning to the mtDNA genome and then used four alternative strategies to obtain sequence variation for mtDNA classification: a) one based on the alignment of reads to the rCRS sequence with Minimap2 (Li, 2018) followed by a variant-calling step with Medaka (https://github.com/nanoporetech/medaka); b) another relying on the reference-based assembly performed by Rebaler (https://github.com/rrwick/Rebaler); c) a third strategy based on *de novo* assembly with Miniasm (Li, 2016) followed by nine rounds of polishing with Racon (Vaser et al., 2017) and a final step with RagTag (Alonge et al., 2019) for scaffolding; and d) the last strategy combining ONT and Illumina reads in a hybrid *de novo* assembly built with Unicycler (Wick et al., 2017) and a final step with RagTag for scaffolding (see supplementary **Figure S2**). Quality of assemblies were assessed with QUAST (Gurevich et al., 2013). For all the alternatives, a VCF file per sample was generated for the haplogroup classification step.

For each sample, three different files were generated for short-read data: BAM and VCF files created through the pipeline previously described, and FASTA files obtained by the VCF-consensus script included in vcftools (Danecek et al., 2011) generated from the VCF files. Of note, an additional filter was applied to the VCF files discarding variants that showed an allele fraction below 0.9, a threshold applied by default in Haplogrep during the haplogroup classification. This enabled the harmonization of the genetic variation considered in FASTA and VCF files.

### mtDNA haplogroup classification

Among the tools available from the literature, we selected 11 that were published in the last three years or that were cited at least 30 times since its description (**Table 1**). Haplogrep, Haplocheck and Phy-Mer have the option of using alternative input files, fostering an evaluation with the alternative supported format files. Some of the tools provide quality scores supporting the mtDNA haplogroup classification. In these cases, only the classification with the best scoring was considered for the analyses. For the haplogroup classification process, the tools that were designed to be run locally, either through an application or via command line, were executed on a 4-core Intel Core i7 CPU at 2.6 GHz and 16 GB of RAM.

**Table 1.**
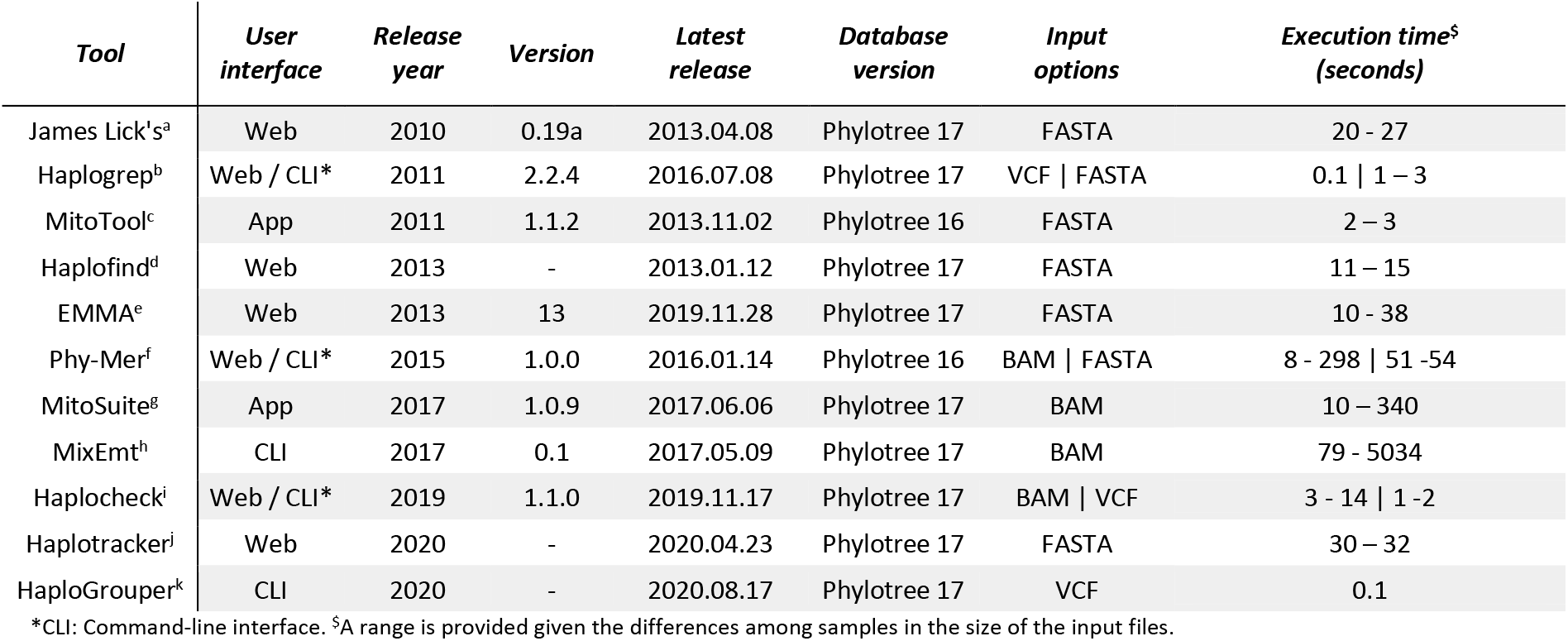
List of software assessed for human mtDNA haplogroup classification. All the tools assessed classify mtDNA sequences according to the PhyloTree nomenclature by using the latest version Build 17. Two of them, MitoTool and Phy-Mer, remain outdated in regards of the PhyloTree built when this study was conducted. a: https://dna.jameslick.com/mthap/; b: (Weissensteiner et al., 2016); c: (Fan & Yao, 2011, 2013); d: (Vianello et al., 2013); e: (Röck et al., 2013); f: (Navarro-Gomez et al., 2015); g: (Ishiya & Ueda, 2017); h: (Vohr et al., 2017); i: (Weissensteiner et al., 2020); j: (Kim et al., 2020); k: (Jagadeesan et al., 2020).

### Statistical analyses

All statistical analyses were performed in R v.3.3.3 (R Core Team 2017). Model fitting was evaluated visually using the ‘DHARMa’ package (Hartig, 2017) (v.0.1.5). Prior to analysis, all predictors were standardized by subtracting the mean and dividing by the standard deviation with the ‘scale’ base function in R.

In order to evaluate the performance of the 11 tools in the classification, we ran generalized linear mixed models (GLMM) with the tools as fixed factor and the classification results (concordance/discordance) as the response variable. This was done separately for WGS and WES data. In the models, we included the sample as a random factor to account for differences introduced by the sequencing runs and the diverse enrichment kits involved. Classification results were transformed to a binary response so that, for each sample, discordance between the consensus haplogroup and the classification result was coded as one, and the concordance as zero. In such binary outcome data, the models may suffer from complete separation when one of the levels of an explanatory variable explains completely the binomial response variable, precluding an optimal fitting of the algorithm (Heinze & Schemper, 2002). This prevents the algorithm from properly fitting the coefficient for this level. To overcome this issue, we fitted the models using the bglmer function of the “blme” v.1.0.4 R package (Chung et al., 2013), which incorporates a similar algorithm to “lmer4” v.1.1.15 R package (Bates et al., 2015) to facilitate the estimation with the incorporation of a prior. Before the analysis, a Pearson’s correlation test was conducted to examine the cross-correlation among the different sequencing parameters measured (i.e., mapped reads, duplicate reads, depth, mean mapping quality, and base quality). Pearson’s correlation test revealed that the proportion of duplicate reads and the depth of coverage were highly correlated (r < 0.8). Therefore, only the mapped reads, mapping quality, and base quality were incorporated as covariates in the model (see supplementary **Table S2**). Finally, to evidence the pairwise differences between the tools, we run a *post-hoc* analysis with the Tukey contrast by using the “multcomp” v.1.4.8 package (Hothorn et al., 2008). Model outputs were visualized using ‘effects’ v.0.9.4 (Fox, 2003) and ‘ggplot2’ v.2.2.1 (Wickham, 2016).

## 3. Results

### Short-read sequencing summary of mitogenomes

The mean (± SD) number of mtDNA reads recovered per sample (n=36) for short-read WGS and WES data were 197,717 ± 98,719 and 9,905 ± 6,808, respectively. For WGS, 100% of the mitogenome was covered at least at 1X. For WES, this percentage decreased to a mean (± SD) of 86.79% ± 27.01 of the recovered mitogenome. The mean (± SD) mapping quality for mapped reads had a value of 59.9 ± 0.01 and the Phred base-quality scores estimated were high for both applications (28.38 ± 0.91 for WGS, and 29.74 ± 0.02 for WES). The average (± SD) depth recovered for WGS was 1,119X ± 433 (range: 554-2,577X), decreasing to 37X ± 20 for WES (range: 11-92X). Besides, while WGS provided a homogeneous depth of coverage profile across the mitogenomes, those recovered by WES showed a highly heterogeneous profile across samples. Interestingly, the region between nucleotide positions 2,000-3,000 was highly enriched in reads from Illumina WES data (**Figure 1**). As for the number of detected variants after filtering, the mean (± SD) depth of coverage per variant call had a value of 958 ± 362 in WGS, decreasing to 40 ± 21 in WES. Out of the 36 samples sequenced, ten showed the equivalent number of variants by WGS and WES. In the remaining samples, the number of variants detected by WES was lower than by WGS: 17 samples missed between 1 and 3 variants, and nine missed more than 30% of the variants (**Table S1**).

**Figure 1.**
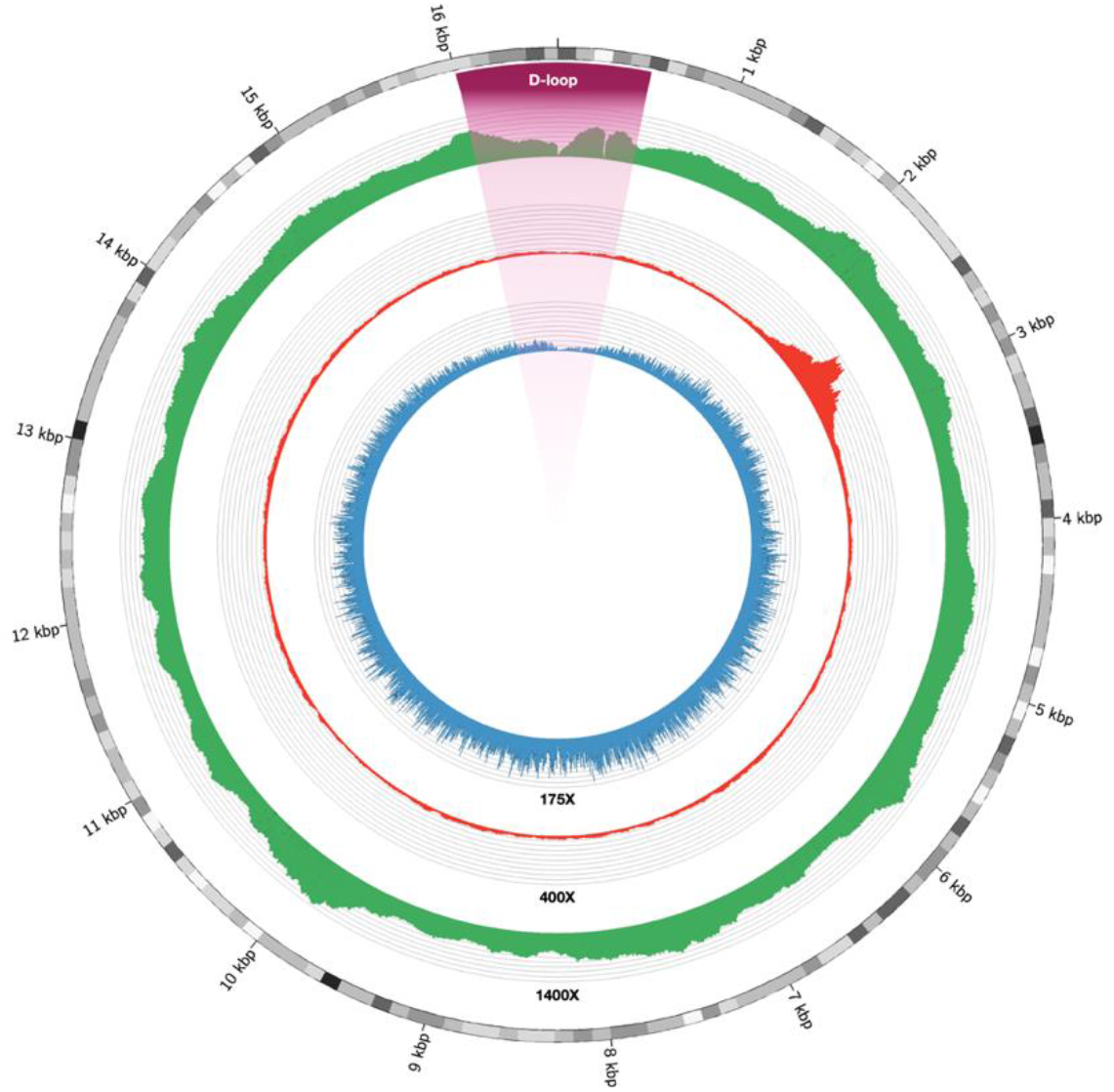
Circos plot of the depth of coverage for short-read and long-read sequencing in the mtDNA of an exemplar sample. Short-read WGS and WES data are colored in green and red, respectively. Long-read WGS data is shown in blue.

### Haplogroup classification based on short-read data

Twenty-eight samples showed >90% concordance in the haplogroup classification among the 11 tools. The remaining showed a mean concordance rate of 69.23% (see supplementary **Table S3)**. In addition, a higher classification accuracy was provided for WGS, reaching an average of 90.08%, while for WES it decreased to an average of 76.98% (see supplementary **Table S4**). On average (± SD), there were more discordances on the WES classifications (2.97 ± 3.92) than for WGS (1.39 ± 1.48). Based on this evidence, we set the WGS-derived haplogroup as the ground truth.

By the classification accuracy, Haplogrep, Haplocheck, EMMA, and James Lick’s mtHap classified all samples correctly based on WGS. For the WES data, only Haplocheck classified all haplogroups correctly, and only when the BAM file was used as input. Concerning the less accurate classifiers, MixEmt showed the lowest accuracy for WGS, only classifying precisely 30.55% of the analyzed samples. MitoTool was the least accurate tool for WES data, yielding an incorrect haplogroup classification in 41.67% of the samples. The execution time required for haplogroup classification was also very different between the tools (**Table 1**). As expected, classifications that relied on BAM inputs had a processing time higher than those running with FASTA and VCF files. Among the tools that support BAM file inputs, MixEmt was the most time-consuming, in which the heterogeneity of processing time among samples was likely due to the different number of mtDNA reads between datasets. In case of Haplocheck, which was designed along with MixEmt to estimate the potential cross-contamination of samples, it required less time per sample for the haplogroup classification.

### Modelling of the haplogroup classification accuracy

Based on these results, we modelled the performance of the 11 tools for short-read mtDNA data classification. The total variance explained by a fixed effects model was 57.69% and 46.19% for WGS and WES data, respectively. The predicted probabilities were then estimated in order to assess the haplogroup classification accuracy of each tool both on WES and WGS data (**Figure 2**). Among all tools, MixEmt (estimate ± SE; 3.55 ± 0.66; *p* < 0.001) was the one with the lowest accuracy in correctly classifying the haplogroup from WGS (80.91% predicted probability of incorrect classification). The rest of the tools showed a negligible probability of misclassifying the haplogroup from WGS (**Figure 2** and **Table S5**). *Post-hoc* tests did not show statistically significant differences among them. For WES, Haplocheck (−4.38, ± 1.99, *p* = 0.03) and MixEmt (−1.13, ± 0.50, *p* = 0.02), both using BAM as the input file, were the tools showing the highest accuracy for haplogroup classification (**Figure 2** and **Table S5**). Haplocheck correctly classified the haplogroup for all samples, while MixEmt incorrectly classified four of the 36 samples analyzed. Despite that, *post-hoc* tests did not show significant differences in the classification performance between Haplocheck and MixEmt. On the contrary, MitoTool (0.97, ± 0.49, *p* = 0.05) showed the lowest accuracy in the haplogroup classification based on WES, for which the predicted probability of incorrectly classifying an haplogroup was ≥25%. Regarding the effect of the covariates, their effect was negligible for WGS data. However, for the WES data, a relatively high effect size was found for the mapped reads (−5.96 ± 3.55; *p* = 0.09) and the mapping quality (1.16 ± 0.39; *p* < 0.001) (**Table S5**). Taken together, these results suggest that the number of reads available and their mapping quality have stronger effects on the probability of inaccurate haplogroup classification based on WES data than based on WGS data.

**Figure 2.**
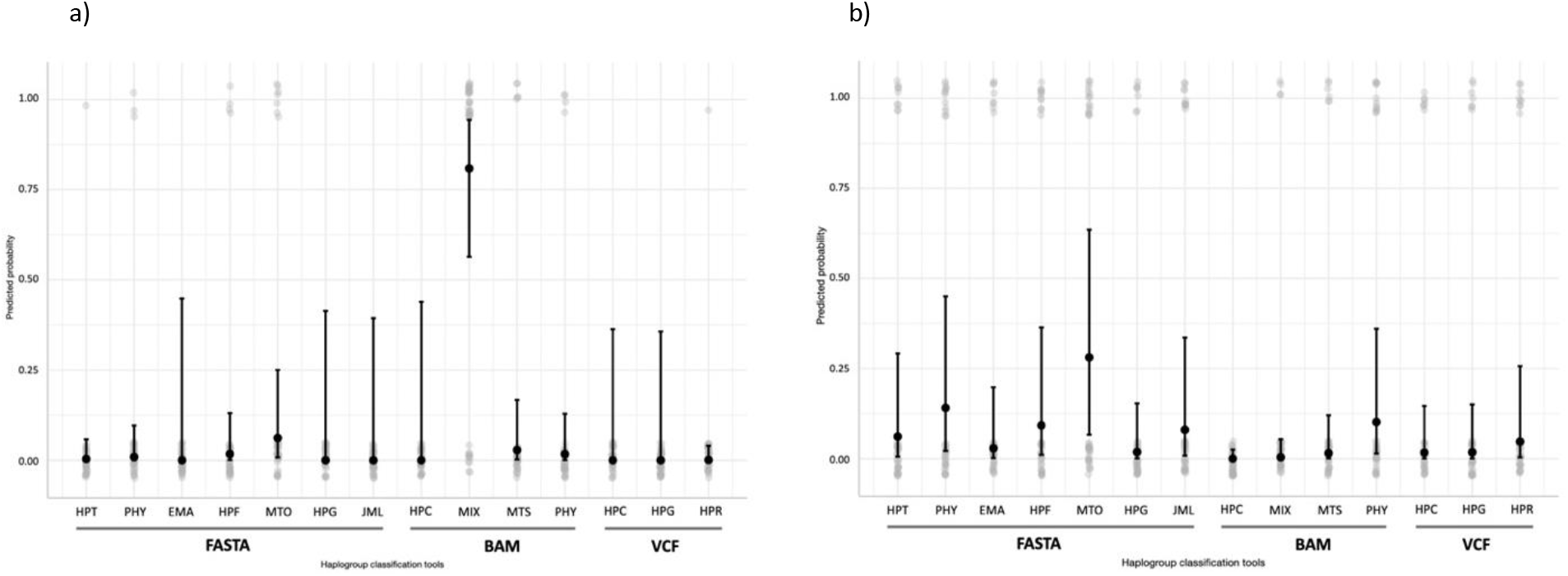
Predicted probability values (and 95% confidence intervals) of the GLMM model estimated for each tool for a) WGS and b) WES datasets. In light grey, the raw data from the haplogroup classification results. **JML**: James Lick’s, **HPG**: Haplogrep, **MTO**: MitoTool, **HPF**: Haplofind, **EMA**: EMMA, **PHY**: Phy-Mer, **MTS**: MitoSuite, **MIX**: MixEmt, **HPC**: Haplocheck, **HPT**: Haplotracker, **HPR**: HaploGrouper.

### Qualitative assessment of haplogroup classification tools

Due to the large number of haplogroup classification tools, a secondary aim of this study was to provide a guidance for researchers to select the most suitable tool for their analyses, not only based on the mtDNA haplogroup classification accuracy but also based on different software features and the usability of the evaluated tools. In order to facilitate the comparison among different tools, a qualitative assessment table is provided with the advantages and limitations of each tool. For this, the following characteristics were considered: haplogroup classification accuracy, computation time for classification, whether or not the latest Phylotree database is used, ability to process cohorts, versatility in the input files supported, user-friendly interface, frequency of tool maintenance, and the presence of others major functions (**Table 2**). Overall, Haplocheck proved to be the most complete tool, achieving the best performance in over 90% of the evaluated features. At the opposite end, Phy-Mer, with more than 50% of the features resulting poorly classified was the tool with the worst performance among all the assessed tools in this study.

**Table 2.**
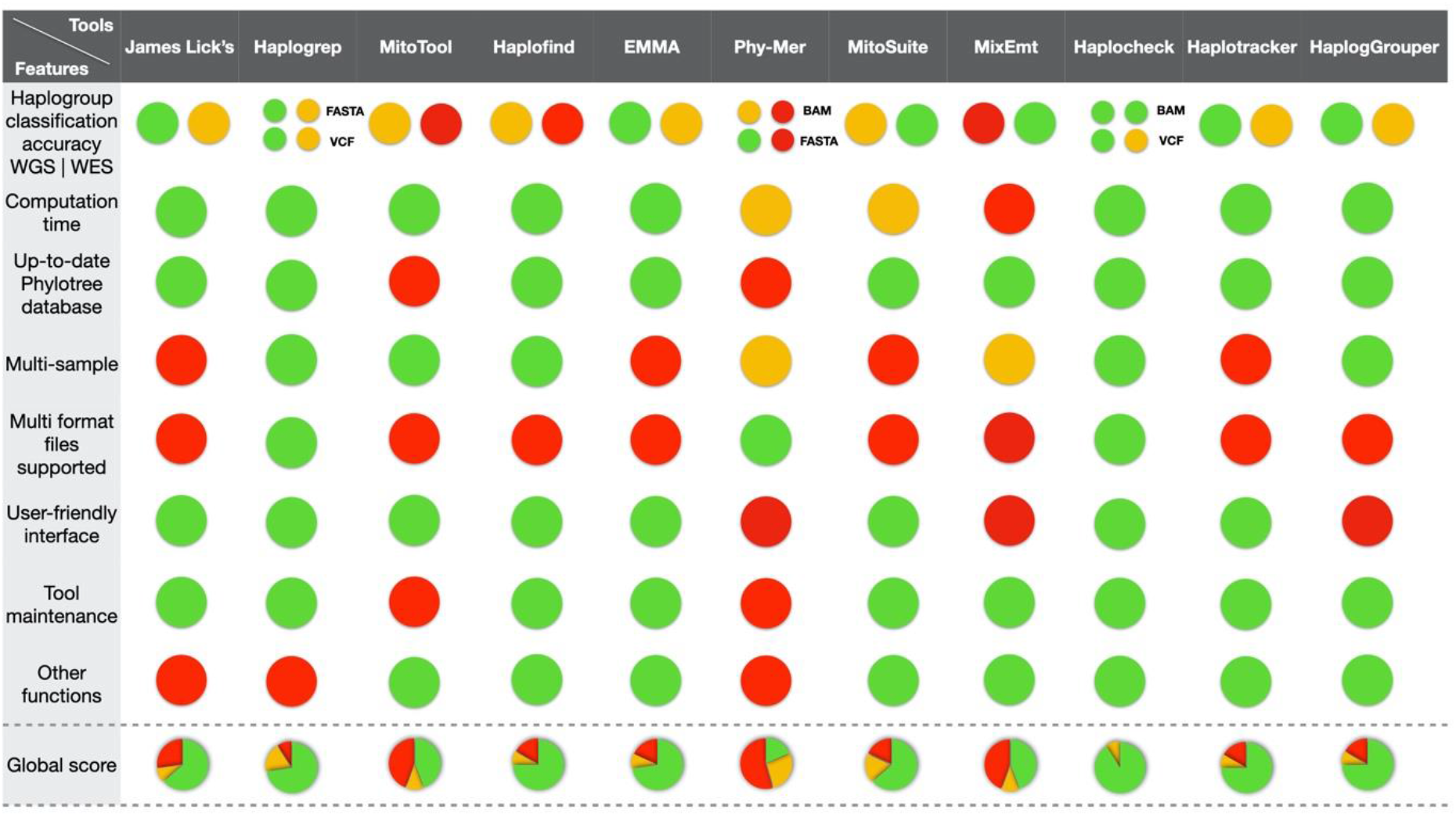
Qualitative assessment of mtDNA haplogroup classification tools. Performance of each tool is evaluated across different features and represented on a color scale based on the level of performance: green for good, orange for fair, and red for low performance. Haplogroup classification accuracy of each tool was categorized into three ranges based on the predicted probabilities by each application: tools with a predicted probability of incorrectly classifying a haplogroup below 1% were represented as good performance, for a fair performance was established a range between 1.01% and 10%, and as low performance tools with a predicted probability higher than 10.01%. For computation time, three intervals were established: tools that classified samples in less than a minute were defined as good performance, from 1 to 5 minutes were categorized as fair performance, and those tools that required more than an hour were represented as low performance. Regarding the PhyloTree database used, it was represented as low performance those tools which are not updated to the latest version of Phylotree, Build 17, and those that allow the latest version as high performance. Multi-sample function was evaluated based on the possibility of cohort analysis. Tools that allow to process these functions were defined as good performance, tools that allow processing several samples by using a loop through a command-line were categorized as fair performance, and tools without this ability were represented as low performance. Based on the file format supported two categories were established: tools that support various input format files categorized as high performance and those that only support one format file that are represented as low performance. The user interface was divided into two categories, tools based on web or desktop applications defined as good performance, and tools developed exclusively for CLI as low performance. The tool maintenance was classified into two classes, tools updated continually or have been recently released, both identified as good performance, and tools that have not been updated during the last years. The last feature is the presence or not of additional functions; tools that have other functions implemented are categorized as good performance. Those tools without more functions were determined as low performance.

### Assessment of ONT data for mtDNA haplogroup classification

The mean (± SD) number of mtDNA reads per sample (n=8) recovered by ONT was 2,812 ± 1,003. The average mtDNA coverage depth was 576X (range: 358-910X). The ONT-recovered mitogenome profile was uniform. However, a decrease in the coverage depth was observed for the D-loop region (positions between 16,024-576) (**Figure 1**). Regarding the quality of the recovered assemblies, the hybrid *de novo* and reference-based assembly strategies reached a similar value of N50, with 16,571 bp and 16,569 bp, respectively. However, for the long-read only *de novo* assembly, this length decreased to 16,407 bp. The consensus overall identity with the rCRS sequence reached a value of 100% for hybrid *de novo* assembly, 99.97% for reference-based strategy, and 88.99% for the long-read only *de novo* assembly strategy. Taking the called variants by short-read WGS as the ground truth, the variant calling strategy for ONT data shared a consensus average (± SD) of 76.41% (± 5.43), the reference-based assembly shared an average of 93.95% (± 5.76), *de novo* assembly shared an average of 89.13% (± 5.56), and the hybrid *de novo* assembly shared an average of 97.35% (± 3.34). In addition, the strategies that only use ONT reads for mtDNA assembly called a larger number of variants that were undetected by short-read WGS. The reference-based assembly strategy called a total of 61 (± 31) variants and *de novo* assembly 158 (± 173). For the variant calling and hybrid *de novo* assembly strategies, the number of novel variants called was lower, decreasing to an average of 3 (± 2) and 1 (± 2), respectively.

Given that Haplocheck was the best performing classifier based on short-read data, we then used it for ONT mtDNA haplogroup classification. The reference-based assembly and the hybrid *de novo* assembly strategies classified all samples correctly. However, both the variant calling and the *de novo* assembly approaches incorrectly classified one out of the eight samples analyzed. Despite that, all samples were classified correctly at the macro-haplogroup level with all strategies (**Table 3**).

**Table 3.**
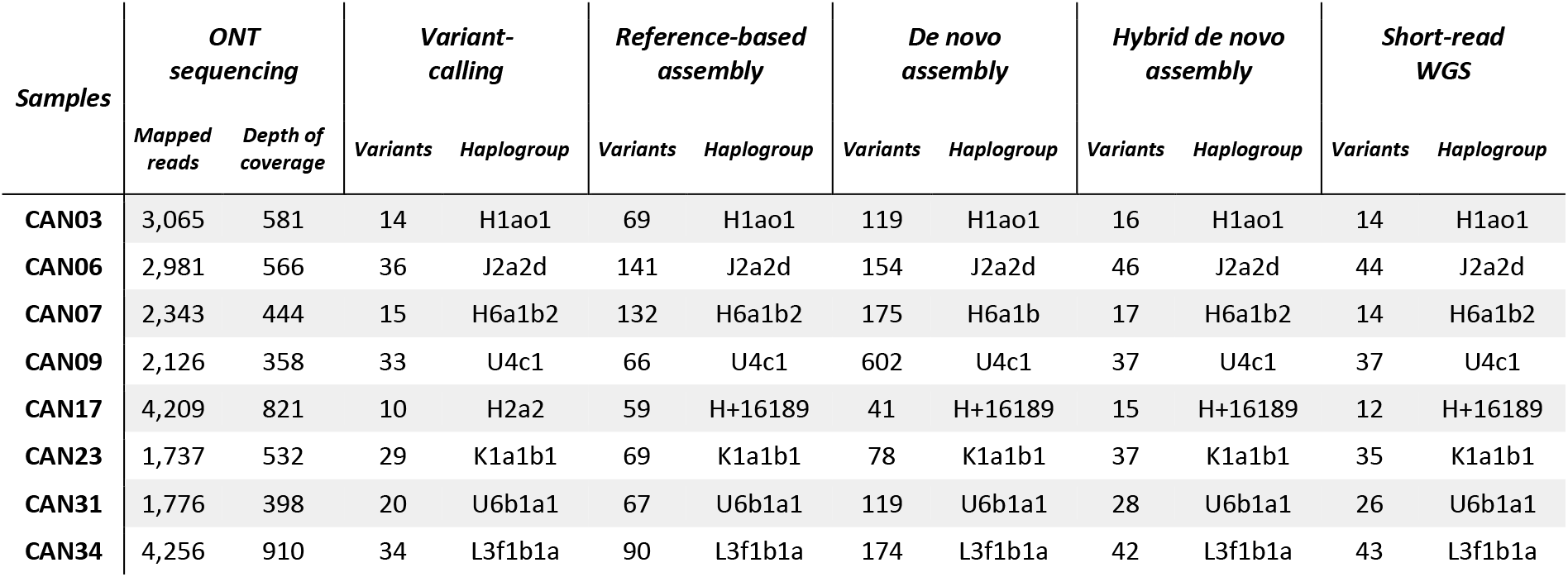
Long-read (ONT) sequencing summary and haplogroup classification results for the samples. The results of the short-read whole-genome sequencing (WGS) mtDNA classification are also shown.

| Samples | ONT sequencing |  | Variant-calling |  | Reference-based assembly |  | De novo assembly |  | Hybrid de novo assembly |  | Short-read WGS |  |
| --- | --- | --- | --- | --- | --- | --- | --- | --- | --- | --- | --- | --- |
|  | Mapped reads | Depth of coverage | Variants | Haplogroup | Variants | Haplogroup | Variants | Haplogroup | Variants | Haplogroup | Variants | Haplogroup |
| CAN03 | 3,065 | 581 | 14 | H1ao1 | 69 | H1ao1 | 119 | H1ao1 | 16 | H1ao1 | 14 | H1ao1 |
| CAN06 | 2,981 | 566 | 36 | J2a2d | 141 | J2a2d | 154 | J2a2d | 46 | J2a2d | 44 | J2a2d |
| CAN07 | 2,343 | 444 | 15 | H6a1b2 | 132 | H6a1b2 | 175 | H6a1b | 17 | H6a1b2 | 14 | H6a1b2 |
| CAN09 | 2,126 | 358 | 33 | U4c1 | 66 | U4c1 | 602 | U4c1 | 37 | U4c1 | 37 | U4c1 |
| CAN17 | 4,209 | 821 | 10 | H2a2 | 59 | H+16189 | 41 | H+16189 | 15 | H+16189 | 12 | H+16189 |
| CAN23 | 1,737 | 532 | 29 | K1a1b1 | 69 | K1a1b1 | 78 | K1a1b1 | 37 | K1a1b1 | 35 | K1a1b1 |
| CAN31 | 1,776 | 398 | 20 | U6b1a1 | 67 | U6b1a1 | 119 | U6b1a1 | 28 | U6b1a1 | 26 | U6b1a1 |
| CAN34 | 4,256 | 910 | 34 | L3f1b1a | 90 | L3f1b1a | 174 | L3f1b1a | 42 | L3f1b1a | 43 | L3f1b1a |

## 4. Discussion

Over the last decade, the number of mtDNA haplogroup classification tools has increased considerably. However, to our best understanding, there was no benchmarking study that assessed the accuracy of the haplogroup classification provided by them until now. This study presents a comparative of the 11 tools that are most widely used. The evaluation was done using empirical HTS data from two of the most widely used applications in human genetics, WES and WGS, in diverse empirical data. Besides, we also evaluated the best performing tool in long noisy ONT reads in a subset of the samples. Our results support that WES offers a suitable solution to allow accurate reconstruction of the mtDNA sequence, although at a lower depth of coverage than WGS. Our results also demonstrated that the most accurate tools for human mtDNA haplogroup classification from short-read WGS data are Haplogrep, Haplocheck, James Lick’s, EMMA, HaploGrouper, and Haplotracker. On the contrary, for short-read WES, Haplocheck and MixEmt were the most accurate. Considering both the accuracy of classification for both approaches and the qualitative assessment made, Haplocheck demonstrated the best performance overall. In fact, using Haplocheck on long noisy ONT reads, we were able to accurately retrieve the mtDNA haplogroup from all assessed samples.

Previous studies have used WES data for human mitogenome reconstruction (Griffin et al., 2014; Patowary et al., 2017; Picardi & Pesole, 2012; Wortmann et al., 2015). Based on our comparisons between WGS and WES datasets, the most striking difference between their classification results was mainly related to the breadth and depth of coverage, both being better for WGS than for WES under our standards. Differences in mtDNA breadth of coverage were less pronounced. However, the mean depth of mtDNA for WGS was around 30 times higher than that obtained by WES. For WGS, the depth reported in diverse studies ranged between 1,200-4,000X (Puttick et al., 2019; Raymond et al., 2018; Watson et al., 2020), fitting with the expected proportion of mtDNA copy number compared to the nuclear DNA, which theoretically differs by 10 to 100 times (Al-Nakeeb et al., 2017; Robin & Wong, 1988). With respect to WES, the observed depth fits in the range described in the literature (Abicht et al., 2018; Diroma et al., 2020; Griffin et al., 2014; Patowary et al., 2017). Attending to the number of variants compared to those detected by WGS, a loss of variants was observed for WES in several samples. This may be explained by the low number of mapped reads of the mtDNA recovered in these samples, causing a shallower mtDNA depth. Despite that, based on the BAM-deduced haplogroup results, we and others (Diroma et al., 2020) have confirmed that WES can be an efficient approach to recover complete human mitogenomes, which allows retrieving the haplogroup with comparable accuracy to that based on WGS.

In general, a loss of accuracy in mtDNA classification was found for WES data in most of the evaluated tools. This could be explained by the lower depth of mitogenomes in WES compared to WGS, as has been described elsewhere (Ishiya & Ueda, 2019; Yin et al., 2019), given that a low depth of coverage will have negative consequences on the performance of the variant calling algorithms (Jennings et al., 2017; Petrackova et al., 2019). mtDNA haplogroup classification relies on a hierarchical algorithm based on the presence or absence of specific diagnostic variants defining the genealogy (Lee et al., 2008). Therefore, missing key variants due to the low depth may have a strong impact on the correct assignment in those tools that only support VCF and/or FASTA files as input. In contrast, the tools that can rely on BAM as an input file, given that this format contains all mapped reads -- including information about the alignment conditions and the complete sequence together with the quality for each base - may facilitate the proper haplogroup classification in sample datasets where certain variant positions are supported by a low number of reads. Hence, the BAM format might be optimal in samples where a low number of mtDNA reads is expected. The use of the BAM file as a possible input file for haplogroup classification has become popular in the last few years (Ishiya & Ueda, 2017; Navarro-Gomez et al., 2015; Vohr et al., 2017; Weissensteiner et al., 2020). With respect to the WES haplogroup results, the tools that support BAM as input files obtained the highest classification accuracy, except Phy-Mer, which was one of the tools performing worst in the classification. Haplocheck and MixEmt reached the highest accuracy among all the tools evaluated. Haplocheck excels in the correct classification of all samples, even with low depth of mtDNA, unlike MixEmt, which resulted in several misclassifications. On the other hand, our results from WES data also showed that the total number of mapped reads, which closely relates to the depth of coverage, has a strong effect on the haplogroup classification, affecting the classification of the samples with a low number of mtDNA reads. However, the effect of this variable was negligible in WGS datasets. Consequently, the total number of mtDNA mapped reads can be considered another key factor to take into account in the mtDNA classification from WES datasets.

Haplogrep, Haplocheck, James Lick’s, EMMA, HaploGrouper, and Haplotracker tools yielded the highest accuracy scores based on WGS data, and their accuracy was independent of the input file format. This finding may be related to the high and uniform depth of WGS throughout the mitogenome, translating in strong support of variants by a high number of reads and, therefore, making more equivalent the information contained in the BAM, VCF, and FASTA files. Haplogrep and Haplocheck, both sharing the underlying algorithm developed first for Haplogrep and now integrated as a module directly in Haplocheck (Weissensteiner et al., 2020), were the ones providing the best ratings for all evaluated features. Despite that, taking into account all the evaluated features, Haplocheck stands out as the most complete haplogroup classification tool for WGS data as it also allows detecting potential sample contaminations based on BAM files. For those users who prefer working with mtDNA alignments in FASTA format and with the sole objective of classifying a limited number of samples by web-based user-friendly tools, EMMA, James Lick’s, HaploGrouper, and Haplotracker are good choices as they all showed similar accuracies as Haplogrep and Haplocheck. EMMA is integrated into the EMPOP platform and stands out as one of the most complete and up-to-date databases of human mtDNA information, containing high quality representations of haplogroups from all over the world based on logical and phylogenetic measures suitable for forensic purposes (Amorim et al., 2019; W. Parson et al., 2014). James Lick’s is one of the first tools released for mitochondrial haplogroup classification that is continually updated to new versions of the PhyloTree database. This tool is widely used among genetic genealogists because it is user friendly and based on a web application. HaploGrouper, the most recently released tool, allows classifying the haplogroups both for mtDNA and the non-recombining portion of the Y-chromosome, being unique in this dual function. However, this tool requires basic bioinformatic skills since it runs from a command-line interface. Finally, Haplotracker has been designed for fragmented DNA samples, such as degraded ones, allowing datasets to be classified using both short reads and complete mtDNA sequences. It runs as a web application with a user-friendly interface that makes it appealing for users without bioinformatics skills.

Long-read sequencing technology allows new genome discoveries leveraging the improvements in genome assemblies and detecting structural variants, among others. Long noisy ONT reads have been applied for studies of the nuclear genome (Beyter et al., 2019; Olson et al., 2020). However, there are few examples of the use of this sequencing technology for mtDNA genome analysis (Franco-Sierra & Díaz-Nieto, 2020; Lindberg et al., 2016). The Achilles’ heel of this emerging sequencing technology, the high error rates, is continuously improving mostly based on pore modifications and the development of basecalling methods. Here we showed that reference-based assembly and hybrid *de novo* assembly strategies provide precise results for haplogroup classification. However, despite the high number of artefactual variants detected using only the long reads for assemblies (reference-based and *de novo)*, these results could be improved using new methods for basecalling and/or genome assembly. Irrespective of that, our results demonstrate that the ONT reads are appropriate for recovering accurate mtDNA haplogroups from WGS data.

## 5. Conclusions

With the advent of the HTS technologies, the number of human mtDNA haplogroup classification tools has increased notably in the last decade. Each new tool released incorporates novel features and different analysis functions, but this has not been always linked to an improvement in the haplogroup classification accuracy. In this study, an evidence-based benchmarking effort was proposed to compare the classification accuracy provided by the most salient tools. We conclude that Haplocheck is the most suitable mtDNA haplogroup estimator for WGS and WES datasets, not only because of its classification accuracy but also because of all the included features and its user-friendly web interface. Regarding third-generation HTS, despite the lower *per* base accuracy currently offered by ONT, we found that it does not hinder a precise human mtDNA classification.

## 6. Acknowledgements

We would like to thank the support from our colleagues from the Teide-HPC Supercomputing facility (http://teidehpc.iter.es/en), which was funded by INP-2011-0063-PCT-430000-ACT (INNPLANTA program) from the Spanish Ministry of Economy and Competitiveness.

## 7. Author Contributions

Conceptualization, VGO, AMB, JMLS, CF; Data curation, VGO, AMB, JMLS, LARR, DJT, RGM; Methodology, VGO, AMB, JMLS, CZT, LARR, ADU, DJT, AIC, RGM; Supervision, CF; Writing—original draft preparation, VGO, AMB, CF; Writing—review and editing, VGO, AMB, JMLS, RGM, CF. All authors have read and agreed to the published version of the manuscript.

