## Supplementary Material for "A benchmarking of human mitochondrial DNA haplogroup classifiers from whole-genome and whole-exome sequence data"

### Supplementary Materials

**Figure S1.** Bioinformatic pipeline for short-read WES and WGS data processing and variant calling of human mitogenomes.

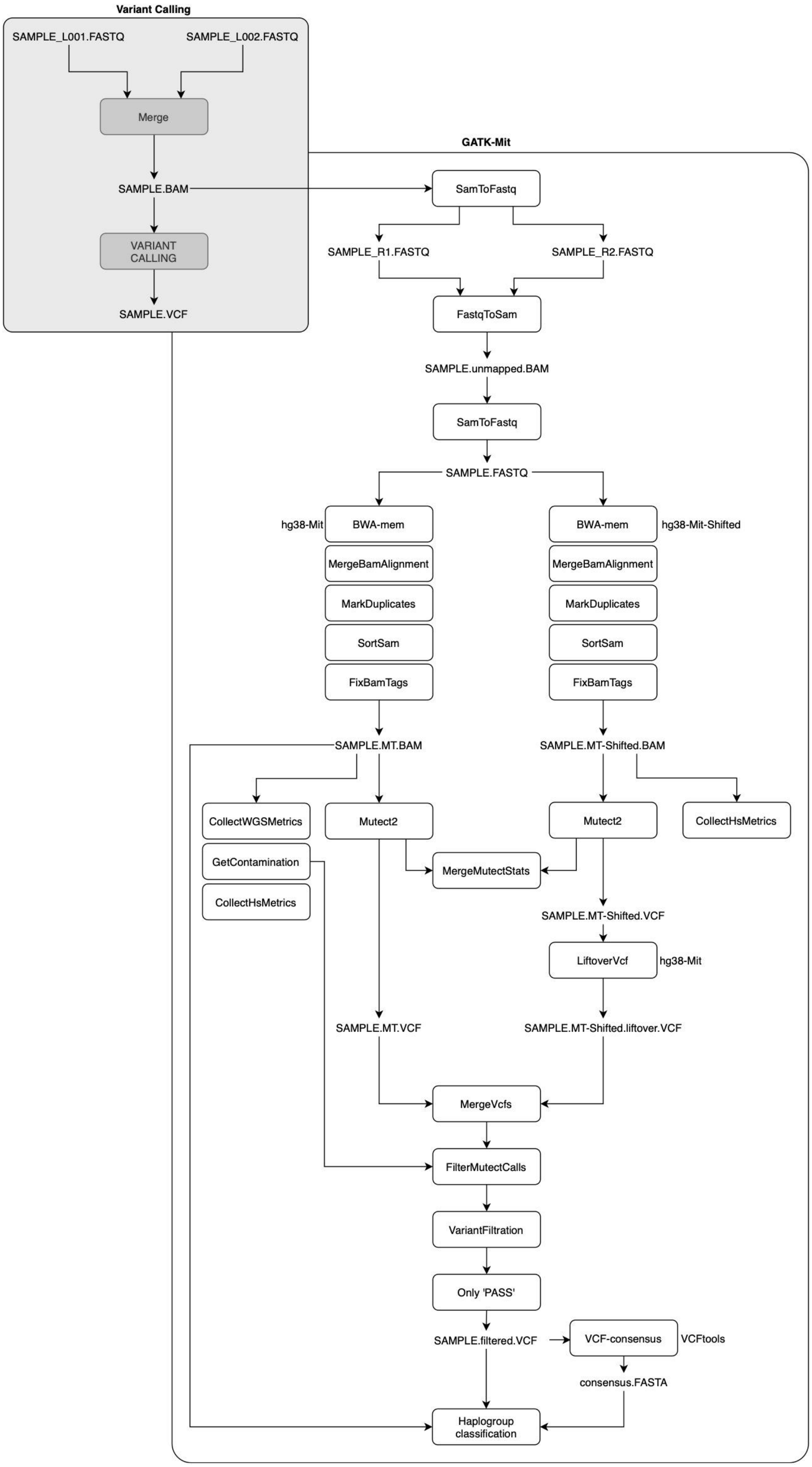

**Figure S2.** Bioinformatic pipeline for processing the long-read ONT data for human mitogenome reconstruction, including the four strategies used in the study: *de novo* assembly using long reads (in green), the hybrid *de novo* assembly using short and long reads (in red), the reference-based assembly using long reads (in blue), and the variant-calling strategy using long reads (in purple).

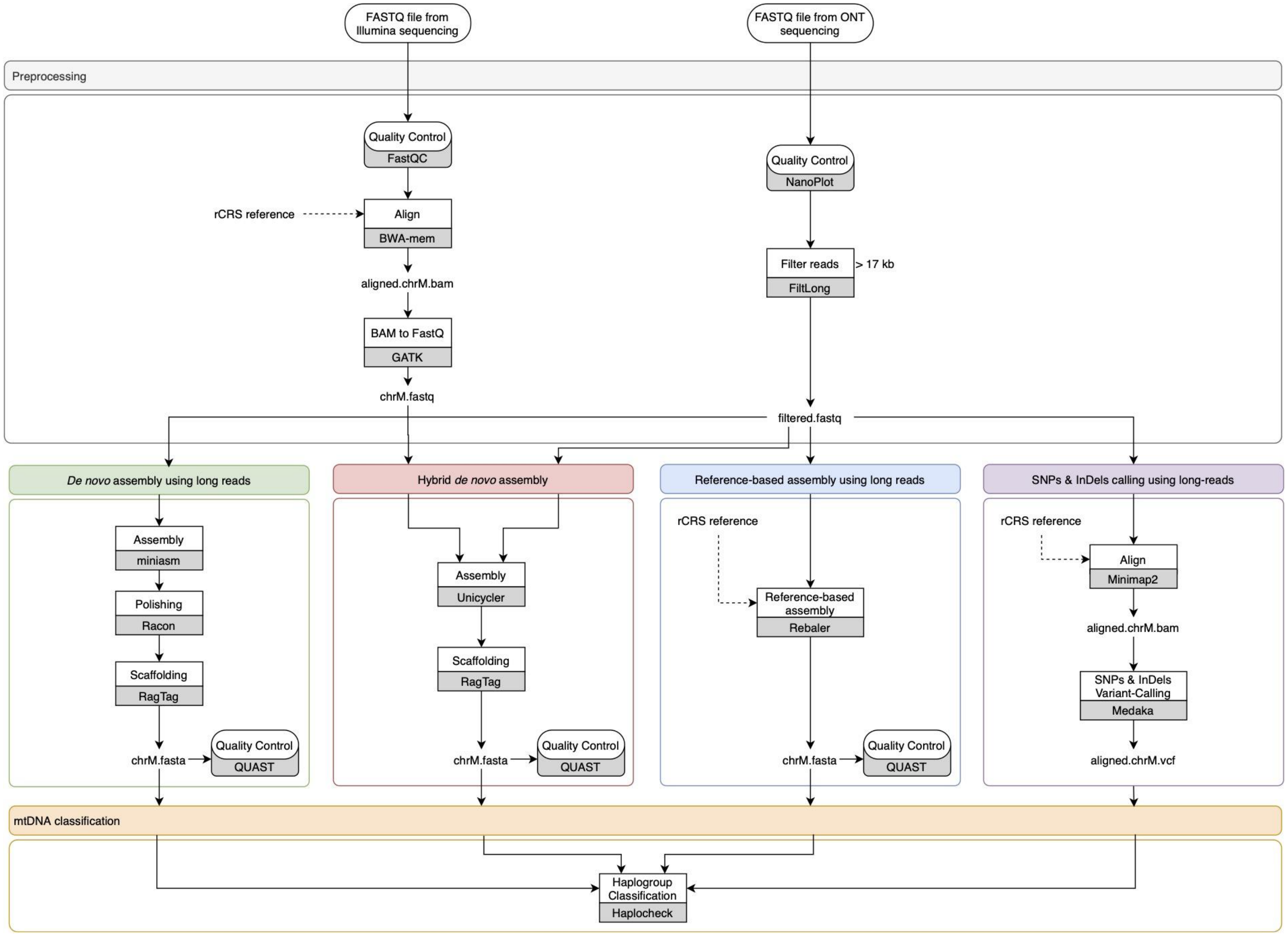

**Table S1.** Short-read sequencing summaries of the samples.

| *Samples* | *Whole-Genome Sequencing* | | | | | | | *Whole-Exome Sequencing* | | | | | | |
|---|---|---|---|---|---|---|---|---|---|---|---|---|---|---|
| | *Protocol* | *Mapped reads* | *Duplicate reads* | *Mapping Quality* | *Base Quality* | *Depth* | *Variants* | *Protocol* | *Mapped reads* | *Duplicate reads* | *Mapping Quality* | *Base Quality* | *Depth* | *Variants* |
| **CAN01** | Nextera-DNA-Prep | 269,488 | 77,240 | 59.98 | 26.00 | 1227 | 42 | Nextera-DNA-Exome | 15,273 | 2,438 | 59.96 | 29.10 | 58 | 42 |
| **CAN02** | Nextera-DNA-Prep | 239,511 | 65,285 | 59.99 | 29.20 | 1466 | 11 | Nextera-DNA-Exome | 13,499 | 1,809 | 59.97 | 29.20 | 53 | 10 |
| **CAN03** | Nextera-DNA-Prep | 225,038 | 63,872 | 59.96 | 28.90 | 1293 | 14 | Nextera-DNA-Exome | 3,299 | 950 | 60.00 | 28.70 | 18 | 6 |
| **CAN04** | Nextera-DNA-Prep | 239,742 | 56,631 | 59.95 | 27.90 | 1589 | 10 | Nextera-DNA-Exome | 16,786 | 2,296 | 59.98 | 29.30 | 66 | 9 |
| **CAN05** | Nextera-DNA-Prep | 140,980 | 27,795 | 59.98 | 28.80 | 974 | 28 | Nextera-DNA-Exome | 9,925 | 1,274 | 59.98 | 29.30 | 39 | 25 |
| **CAN06** | Nextera-DNA-Prep | 209,422 | 86,736 | 59.98 | 28.60 | 977 | 44 | Nextera-DNA-Exome | 15,609 | 2,394 | 59.98 | 29.40 | 60 | 44 |
| **CAN07** | Nextera-DNA-Prep | 202,279 | 53,246 | 59.98 | 29.20 | 676 | 14 | Nextera-DNA-Exome | 17,727 | 3,247 | 59.98 | 29.20 | 66 | 14 |
| **CAN08** | Nextera-DNA-Prep | 102,834 | 22,765 | 59.97 | 28.70 | 661 | 41 | Nextera-DNA-Exome | 2,456 | 702 | 60.00 | 29.20 | 14 | 8 |
| **CAN09** | Nextera-DNA-Prep | 288,083 | 92,888 | 59.99 | 29.40 | 880 | 37 | Nextera-DNA-Exome | 36,117 | 15,507 | 59.99 | 29.50 | 93 | 34 |
| **CAN10** | Nextera-DNA-Prep | 107,424 | 17,424 | 59.98 | 28.80 | 760 | 26 | Nextera-DNA-Exome | 6,501 | 1,082 | 59.97 | 28.70 | 25 | 26 |
| **CAN11** | Nextera-DNA-Prep | 486,352 | 184,988 | 59.97 | 28.60 | 2577 | 25 | Nextera-DNA-Exome | 5,452 | 1,783 | 60.00 | 29.10 | 29 | 17 |
| **CAN12** | Illumina-DNA-Prep | 96,563 | 15,042 | 59.95 | 28.10 | 724 | 25 | Nextera-DNA-Exome | 5,297 | 850 | 59.95 | 28.70 | 20 | 22 |
| **CAN13** | Nextera-DNA-Prep | 298,707 | 84,386 | 59.98 | 29.00 | 1822 | 25 | Nextera-DNA-Exome | 23,503 | 3,853 | 59.99 | 29.30 | 89 | 25 |
| **CAN14** | Nextera-DNA-Prep | 160,800 | 31,404 | 59.98 | 28.60 | 1130 | 12 | Nextera-DNA-Exome | 8,746 | 3,222 | 60.00 | 29.40 | 25 | 12 |
| **CAN15** | Illumina-DNA-Prep | 86,288 | 12,342 | 59.94 | 27.90 | 656 | 34 | Nextera-DNA-Exome | 6,208 | 1,168 | 59.98 | 29.50 | 23 | 31 |
| **CAN16** | Nextera-DNA-Prep | 145,985 | 52,881 | 59.95 | 27.00 | 786 | 33 | Nextera-DNA-Exome | 5,211 | 1,964 | 60.00 | 29.10 | 23 | 20 |
| **CAN17** | Nextera-DNA-Prep | 287,742 | 80,049 | 59.96 | 28.40 | 1691 | 12 | Nextera-DNA-Exome | 5,532 | 1,705 | 60.00 | 28.80 | 30 | 9 |
| **CAN18** | Nextera-DNA-Prep | 186,339 | 78,863 | 59.99 | 28.90 | 874 | 34 | Nextera-DNA-Exome | 5,056 | 2,026 | 60.00 | 29.30 | 22 | 11 |
| **CAN19** | Nextera-DNA-Prep | 158,760 | 34,517 | 59.97 | 28.60 | 1021 | 13 | Nextera-DNA-Exome | 2,345 | 698 | 59.98 | 28.10 | 13 | 4 |
| **CAN20** | Nextera-DNA-Prep | 198,085 | 48,750 | 59.99 | 29.20 | 1284 | 13 | Nextera-DNA-Exome | 9,765 | 1,655 | 59.98 | 36.80 | 37 | 12 |
| **CAN21** | Nextera-DNA-Prep | 183,878 | 37,268 | 59.96 | 28.30 | 1282 | 17 | Nextera-DNA-Exome | 6,736 | 965 | 59.99 | 36.60 | 26 | 17 |
| **CAN22** | Nextera-DNA-Prep | 466,404 | 196,793 | 59.98 | 29.10 | 1823 | 7 | Nextera-DNA-Exome | 3,911 | 1,212 | 59.99 | 29.20 | 22 | 1 |
| **CAN23** | Nextera-DNA-Prep | 149,371 | 32,434 | 59.99 | 28.60 | 1002 | 35 | Nextera-DNA-Exome | 9,810 | 1,422 | 59.99 | 29.20 | 38 | 35 |
| **CAN24** | Nextera-DNA-Prep | 75,746 | 10,746 | 59.97 | 28.70 | 554 | 34 | Nextera-DNA-Exome | 6,809 | 1,010 | 59.95 | 29.20 | 26 | 32 |
| **CAN25** | Nextera-DNA-Prep | 130,389 | 23,037 | 59.97 | 28.40 | 935 | 39 | Nextera-DNA-Exome | 6,707 | 1,184 | 59.99 | 29.30 | 25 | 37 |
| **CAN26** | Illumina-DNA-Prep | 145,401 | 30,564 | 59.96 | 28.40 | 1024 | 35 | Nextera-DNA-Exome | 10,105 | 1,900 | 59.99 | 29.50 | 37 | 32 |
| **CAN27** | Illumina-DNA-Prep | 106,440 | 16,750 | 59.94 | 28.00 | 796 | 39 | Nextera-DNA-Exome | 6,087 | 996 | 60.00 | 29.40 | 23 | 38 |
| **CAN28** | Nextera-DNA-Prep | 341,321 | 102,609 | 59.97 | 25.80 | 1594 | 10 | Nextera-DNA-Exome | 13,280 | 2,376 | 59.98 | 29.20 | 50 | 9 |
| **CAN29** | Nextera-DNA-Prep | 351,160 | 108,741 | 59.97 | 25.80 | 1618 | 15 | Nextera-DNA-Exome | 17,020 | 5,915 | 59.99 | 29.60 | 50 | 15 |
| **CAN30** | Nextera-DNA-Prep | 189,623 | 78,268 | 59.97 | 27.90 | 973 | 38 | Nextera-DNA-Exome | 4,347 | 1,745 | 60.00 | 29.20 | 19 | 11 |
| **CAN31** | Nextera-DNA-Prep | 143,579 | 32,325 | 59.98 | 29.20 | 950 | 26 | Nextera-DNA-Exome | 10,019 | 2,159 | 59.93 | 28.70 | 36 | 25 |
| **CAN32** | Nextera-DNA-Prep | 116,947 | 21,154 | 59.98 | 28.60 | 824 | 26 | Nextera-DNA-Exome | 7,947 | 1,362 | 59.98 | 29.10 | 30 | 25 |
| **CAN33** | Nextera-DNA-Prep | 138,396 | 38,670 | 59.99 | 29.30 | 649 | 31 | Nextera-DNA-Exome | 4,038 | 1,204 | 60.00 | 29.10 | 23 | 17 |
| **CAN34** | Nextera-DNA-Prep | 130,484 | 23,846 | 59.98 | 28.80 | 913 | 43 | Nextera-DNA-Exome | 14,569 | 2,380 | 59.98 | 29.00 | 55 | 42 |
| **CAN35** | Illumina-DNA-Prep | 119,570 | 22,308 | 59.96 | 28.50 | 867 | 40 | Nextera-DNA-Exome | 5,709 | 732 | 59.97 | 35.10 | 23 | 38 |
| **CAN36** | Illumina-DNA-Prep | 198,697 | 40,465 | 59.95 | 28.30 | 1402 | 26 | Nextera-DNA-Exome | 15,194 | 2,908 | 59.99 | 29.50 | 56 | 26 |

**Table S2.** Multicollinearity test results (Pearson correlation coefficients) based on cross-correlation among the selected parameters estimated from the BAM files for short-read A) WGS and B) WES datasets. The statistical significance is indicated by asterisks: * $p \leq 0.05$ | ** $p \leq 0.01$ | *** $p \leq 0.001$

a)

| *R square WGS* | *Mapped reads* | *Duplicate reads* | *Mapping Quality* | *Base quality* | *Coverage* |
|---|---|---|---|---|---|
| ***Mapped reads*** | 1.000 | 0.959*** | 0.190 | -0.197 | 0.877*** |
| ***Duplicates reads*** | 0.959*** | 1.000 | 0.224 | -0.135 | 0.772*** |
| ***Mapping quality*** | 0.190 | 0.224 | 1.000 | 0.382* | 0.018 |
| ***Base quality*** | -0.197 | -0.135 | 0.382* | 1.000 | -0.172 |
| ***Depth*** | 0.877*** | 0.772*** | 0.018 | -0.172 | 1.000 |

b)

| *R square WES* | *Mapped reads* | *Duplicate reads* | *Mapping Quality* | *Base quality* | *Coverage* |
|---|---|---|---|---|---|
| ***Mapped reads*** | 1.000 | 0.837*** | -0.107 | -0.040 | 0.960*** |
| ***Duplicates reads*** | 0.837*** | 1.000 | 0.095 | -0.076 | 0.677*** |
| ***Mapping quality*** | -0.107 | 0.095 | 1.000 | -0.006 | -0.113 |
| ***Base quality*** | -0.040 | -0.076 | -0.006 | 1.000 | -0.064 |
| ***Depth*** | 0.960*** | 0.677*** | -0.113 | -0.064 | 1.000 |

**Table S3.** Mitochondrial DNA haplogroup consensus and concordance rate among the 11 tools based on short-read WGS for the samples assessed.

| *Samples* | *Haplogroup* | *Concordance rate* |
|---|---|---|
| **CAN01** | J2a2d | 100.00 |
| **CAN02** | H7 | 100.00 |
| **CAN03** | H1ao1 | 100.00 |
| **CAN04** | H1t | 92.31 |
| **CAN05** | U6b1a | 100.00 |
| **CAN06** | J2a2d | 92.31 |
| **CAN07** | H6a1b2 | 100.00 |
| **CAN08** | L3d1b3a | 92.31 |
| **CAN09** | U4c1 | 69.23 |
| **CAN10** | U6b1a | 92.31 |
| **CAN11** | U6b1a | 92.31 |
| **CAN12** | U6b1a | 100.00 |
| **CAN13** | U6b1a | 100.00 |
| **CAN14** | H1e1a | 92.31 |
| **CAN15** | U4c1 | 69.23 |
| **CAN16** | T1a | 92.31 |
| **CAN17** | H1+16189 | 69.23 |
| **CAN18** | K1a4a1 | 100.00 |
| **CAN19** | H1e1a | 100.00 |
| **CAN20** | H1au | 92.31 |
| **CAN21** | HV+16311 | 69.23 |
| **CAN22** | H | 69.23 |
| **CAN23** | K1a1b1 | 92.31 |
| **CAN24** | K1a14 | 92.31 |
| **CAN25** | J1c2c1 | 92.31 |
| **CAN26** | J1c2 | 92.31 |
| **CAN27** | T2c1d+152 | 69.23 |
| **CAN28** | H1cf | 69.23 |
| **CAN29** | H6a1a | 92.31 |
| **CAN30** | U5b2b3a | 92.31 |
| **CAN31** | U6b1a1 | 100.00 |
| **CAN32** | U6b1a | 92.31 |
| **CAN33** | X3a | 92.31 |
| **CAN34** | L3f1b1a | 92.31 |
| **CAN35** | L3b1a+@16124 | 69.23 |
| **CAN36** | U6b1a1 | 100.00 |

**Table S4.** Per sample mtDNA haplogroup classifications for the 11 tools based on short-read A) WGS and B) WES datasets.

a)

| *Samples* | *Haplogrep VCF* | *Haplogrep FASTA* | *Haplocheck BAM* | *Haplocheck VCF* | *Haplotracker FASTA* | *HaploGrouper VCF* | *Phy-Mer FASTA* | *Phy-Mer BAM* | *EMMA FASTA* | *MitoSuite BAM* | *Haplofind FASTA* | *MixEmt BAM* | *MitoTool FASTA* | *James Lick's FASTA* |
|---|---|---|---|---|---|---|---|---|---|---|---|---|---|---|
| **CAN01** | J2a2d | J2a2d | J2a2d | J2a2d | J2a2d | J2a2d | J2a2d | J2a2d | J2a2d | J2a2d | J2a2d | J2a2a | J2a2d | J2a2d |
| **CAN02** | H7 | H7 | H7 | H7 | H7 | H7 | H7 | H7 | H7 | H27e | H7 | H22 | H7 | H7 |
| **CAN03** | H1ao1 | H1ao1 | H1ao1 | H1ao1 | H1ao1 | H1ao1 | H1ao1 | H1ao1 | H1ao1 | H1ao1 | H1ao1 | H1ao1 | H1ao1 | H1ao1 |
| **CAN04** | H1t | H1t | H1t | H1t | H1t | H1t | H1t | H1t | H1t | H1t | H1t | H1t2 | H1t | H1t |
| **CAN05** | U6b1a | U6b1a | U6b1a | U6b1a | U6b1a | U6b1a | U6b1a | U6b1a | U6b1a | U6b1a | U6b1a | U6b1a | U6b1a | U6b1a |
| **CAN06** | J2a2d | J2a2d | J2a2d | J2a2d | J2a2d | J2a2d | J2a2d | J2a2d | J2a2d | J2a2d | J2a2d | J2a2a | J2a2d | J2a2d |
| **CAN07** | H6a1b2 | H6a1b2 | H6a1b2 | H6a1b2 | H6a1b2 | H6a1b2 | H6a1b2 | H6a1b2 | H6a1b2 | H6a1b2 | H6a1b2 | H6a1b2b | H6a1b2 | H6a1b2 |
| **CAN08** | L3d1b3a | L3d1b3a | L3d1b3a | L3d1b3a | L3d1b3a | L3d1b3a | L3d1b3a | L3d1b3a | L3d1b3a | L3d1b3a | L3d1b3a | L3d4a | L3d1b3a | L3d1b3a |
| **CAN09** | U4c1 | U4c1 | U4c1 | U4c1 | U4c1 | U4c1 | U4c1a | U4c1a | U4c1 | U4c1 | U4c1 | U4a2g | U4c1a | U4c1 |
| **CAN10** | U6b1a | U6b1a | U6b1a | U6b1a | U6b1a | U6b1a | U6b1a | U6b1a | U6b1a | U6b1a | U6b1a | U6b1a1 | U6b1a | U6b1a |
| **CAN11** | U6b1a | U6b1a | U6b1a | U6b1a | U6b1a | U6b1a | U6b1a | U6b1a | U6b1a | U6b1a | U6b1a | U6b1a1 | U6b1a | U6b1a |
| **CAN12** | U6b1a | U6b1a | U6b1a | U6b1a | U6b1a | U6b1a | U6b1a | U6b1a | U6b1a | U6b1a | U6b1a | U6b1a | U6b1a | U6b1a |
| **CAN13** | U6b1a | U6b1a | U6b1a | U6b1a | U6b1a | U6b1a | U6b1a | U6b1a | U6b1a | U6b1a | U6b1a | U6b1a | U6b1a | U6b1a |
| **CAN14** | H1e1a | H1e1a | H1e1a | H1e1a | H1e1a | H1e1a | H1e1a | H1e1a | H1e1a | H1e1a | H1e1a | H1a4 | H1e1a | H1e1a |
| **CAN15** | U4c1 | U4c1 | U4c1 | U4c1 | U4c1 | U4c1 | U4c1a | U4c1a | U4c1 | U4c1 | U4c1 | U4c1a | U4c1a | U4c1 |
| **CAN16** | T1a | T1a | T1a | T1a | T1a | T1a | T1a | T1a | T1a | T1a | T1a | T1a1b1 | T1a | T1a |
| **CAN17** | H1+16189 | H1+16189 | H1+16189 | H1+16189 | H1+16189 | H1+16189 | H1-16189 | H1-16189 | H1+16189 | B | H1 | B4a1a1ab | H1 | H1+16189 |
| **CAN18** | K1a4a1 | K1a4a1 | K1a4a1 | K1a4a1 | K1a4a1 | K1a4a1 | K1a4a1 | K1a4a1 | K1a4a1 | K1a4a1 | K1a4a1 | K1a4a1 | K1a4a1 | K1a4a1 |
| **CAN19** | H1e1a | H1e1a | H1e1a | H1e1a | H1e1a | H1e1a | H1e1a | H1e1a | H1e1a | H1e1a | H1e1a | H1e1a1 | H1e1a | H1e1a |
| **CAN20** | H1au | H1au | H1au | H1au | H1au | H1au | H1au | H1au | H1au | H1au | H1au | H28 | H1au | H1au |
| **CAN21** | HV+16311 | HV+16311 | H+16311 | HV+16311 | HV+16311 | HV+16311 | HV-16311 | HV-16311 | HV+16311 | B | HV | R0a'b | HV | HV+16311 |
| **CAN22** | H | H | H | H | H | H77 | H | H-16291 | H | H27e | H | H3b | H | H |
| **CAN23** | K1a1b1 | K1a1b1 | K1a1b1 | K1a1b1 | K1a1b1 | K1a1b1 | K1a1b1 | K1a1b1 | K1a1b1 | K1a1b1 | K1a1b1 | K1a1b1b | K1a1b1 | K1a1b1 |
| **CAN24** | K1a14 | K1a14 | K1a14 | K1a14 | K1a14 | K1a14 | K1a14 | K1a14 | K1a14 | K1a14 | K1a14 | K1a5a | K1a14 | K1a14 |
| **CAN25** | J1c2c1 | J1c2c1 | J1c2c1 | J1c2c1 | J1c2c1 | J1c2c1 | J1c2c1 | J1c2c1 | J1c2c1 | J1c2c1 | J1c2c1 | J1c2h | J1c2c1 | J1c2c1 |
| **CAN26** | J1c2 | J1c2 | J1c2 | J1c2 | J1c2 | J1c2 | J1c2 | J1c2 | J1c2 | J1c2 | J1c2 | J1c4c | J1c2 | J1c2 |
| **CAN27** | T2c1d+152 | T2c1d+152 | T2c1d+152 | T2c1d+152 | T2c1d+152 | T2c1d+152 | T2c1d-152 | T2c1d-152 | T2c1d+152 | T2c1d2 | T2c1d2 | T2c1d[1] | T2c1d | T2c1d+152 |
| **CAN28** | H1cf | H1cf | H1cf | H1cf | H1cf | H1cf | H1 | H1 | H1cf | H1cf | H1cf | H3ac | H1 | H1cf |
| **CAN29** | H6a1a | H6a1a | H6a1a | H6a1a | H6a1a | H6a1a | H6a1a | H6a1a | H6a1a | H6a1a | H6a1a | H6a1a2b1 | H6a1a | H6a1a |
| **CAN30** | U5b2b3a | U5b2b3a | U5b2b3a | U5b2b3a | U5b2b3a | U5b2b3a | U5b2b3a | U5b2b3a | U5b2b3a | U5b2b3a | U5b2b3a | U5b2b | U5b2b3a | U5b2b3a |
| **CAN31** | U6b1a1 | U6b1a1 | U6b1a1 | U6b1a1 | U6b1a1 | U6b1a1 | U6b1a1 | U6b1a1 | U6b1a1 | U6b1a1 | U6b1a1 | U6b1a | U6b1a1 | U6b1a1 |
| **CAN32** | U6b1a | U6b1a | U6b1a | U6b1a | U6b1a | U6b1a | U6b1a | U6b1a | U6b1a | U6b1a | U6b1a | U6b1a1 | U6b1a | U6b1a |
| **CAN33** | X3a | X3a | X3a | X3a | X3a | X3a | X3a | X3a | X3a | X3a | X3a | U4b1a1a1 | X3a | X3a |
| **CAN34** | L3f1b1a | L3f1b1a | L3f1b1a | L3f1b1a | L3f1b | L3f1b1a | L3f1b1a | L3f1b1a | L3f1b1a | L3f1b1a | L3f1b1a | L3f1b1a | L3f1b1a | L3f1b1a |
| **CAN35** | L3b1a+@16124 | L3b1a+@16124 | L3b1a+@16124 | L3b1a+@16124 | L3b1a+@16124 | L3b1a+@16124 | L3b1a-@16124 | L3b1a-@16124 | L3b1a+@16124 | L3b1a11 | L3b1a | L3b1a3 | L3b1a | L3b1a-@16124 |
| **CAN36** | U6b1a1 | U6b1a1 | U6b1a1 | U6b1a1 | U6b1a1 | U6b1a1 | U6b1a1 | U6b1a1 | U6b1a1 | U6b1a1 | U6b1a1 | U6b1a1 | U6b1a1 | U6b1a1 |

b)

| *Samples* | *Haplogrep VCF* | *Haplogrep FASTA* | *Haplocheck BAM* | *Haplocheck VCF* | *Haplotracker FASTA* | *HaploGrouper VCF* | *Phy-Mer FASTA* | *Phy-Mer BAM* | *EMMA FASTA* | *MitoSuite BAM* | *Haplofind FASTA* | *MixEmt BAM* | *MitoTool FASTA* | *James Lick's FASTA* |
|---|---|---|---|---|---|---|---|---|---|---|---|---|---|---|
| **CAN01** | J2a2d | J2a2d | J2a2d | J2a2d | J2a2d | J2a2d | J2a2d | J2a2d | J2a2d | J2a2d | J2a2d | J2a2d | J2a2d | J2a2d |
| **CAN02** | H7 | H7 | H7 | H7 | H7 | H7 | H7 | H7 | H7 | H27e | H7 | H7 | H7 | H7 |
| **CAN03** | H1ao | H1ao | H1ao1 | H1ao | H2a2a1 | H2a2a1 | H2a2a1 | H1x | H2a2a1 | H1ao1 | H2a2a1 | H1ao1 | H2a2a1 | H2a2a1 |
| **CAN04** | H1t | H1t | H1t | H1t | H1t | H1t | H1t | H1t | H1t | H1t | H1t | H1t | H1t | H1t |
| **CAN05** | U6b1a | U6b1a | U6b1a | U6b1a | U6b1a | U6b1a | U6b1a | U6b1a | U6b1a | U6b1a | U6b1a | U6b1a | U6b1a | U6b1a |
| **CAN06** | J2a2d | J2a2d | J2a2d | J2a2d | J2a2d | J2a2d | J2a2d | J2a2d | J2a2d | J2a2d | J2a2d | J2a2d | J2a2d | J2a2d |
| **CAN07** | H6a1b2 | H6a1b2 | H6a1b2 | H6a1b2 | H6a1b2 | H6a1b2 | H6a1b2 | H6a1b2 | H6a1b2 | H6a1b2 | H6a1b2 | H6a1b2 | H6a1b2 | H6a1b2 |
| **CAN08** | L3d | L3d | L3d1b3a | L3d | H2a2a1 | H2a2a1 | H2a2a1 | L3d1b1a | H2a2a1 | L3d1b3a | H2a2a1 | L3d1b3a | H2a2a1 | H2a2a1 |
| **CAN09** | U4c1 | U4c1 | U4c1 | U4c1 | U4c1 | U4c1 | U4c1a | U4c1a | U4c1 | U4c1 | U4c1 | U4c1 | U4c1a | U4c1 |
| **CAN10** | U6b1a | U6b1a | U6b1a | U6b1a | U6b1a | U6b1a | U6b1a | U6b1a | U6b1a | U6b1a | U6b1a | U6b1a | U6b1a | U6b1a |
| **CAN11** | U6b1a | U6b1a | U6b1a | U6b1a | U6+16311 | U6a'b'd T16311C | U6a'b'd-16311 | U6b1a | U6b1a | U6b1a | U6b1a | U6b1a | U6a'b'd | U6a'b'd |
| **CAN12** | U6b1a | U6b1a | U6b1a | U6b1a | U6b1a | U6b1a | U6b1a | U6b1a | U6b1a | U6b1a | U6b1a | U6b1a | U6b1a | U6b1a |
| **CAN13** | U6b1a | U6b1a | U6b1a | U6b1a | U6b1a | U6b1a | U6b1a | U6b1a | U6b1a | U6b1a | U6b1a | U6b1a | U6b1a | U6b1a |
| **CAN14** | H1e1a | H1e1a | H1e1a | H1e1a | H1e1a | H1e1a | H1e1a | H1e1a | H1e1a | H1e1a | H1e1a | H1e1a | H1e1a | H1e1a |
| **CAN15** | U4c1 | U4c1 | U4c1 | U4c1 | U4c1 | U4c1 | U4c1a | U4a2c | U4c1 | U4c1 | U4c1 | U4c1 | U4c1a | U4c1 |
| **CAN16** | T1a | T1a | T1a | T1a | T1a | T1a | T1a | T1a6 | T1a | T1a | T1a | T1a | T1a | T1 |
| **CAN17** | H2 | H2 | H1+16189 | H2 | H1a | H1a | H1a | H1a2 | H2 | B | H1a | H1 | H1 | H2 |
| **CAN18** | K1a4 | K1a4 | K1a4a1 | K1a4 | H2a2 | H2a2 | H2a2a | K1a4a1 | H2a2a | K1a4a1 | H2a2a | K1a4a1 | H2a2 | H2a2a |
| **CAN19** | H2a | H2a | H1e1a | H2a | H2a | H2a | H2a | H1e1b1a | H2a | H1e1a | H2a | H1e1a | H2a | H2a |
| **CAN20** | H1au | H1au | H1au | H1au | H1au | H1au | H1au | H1-16311 | H1au | H1au | H1au | H1au | H1au | H1au |
| **CAN21** | HV+16311 | HV+16311 | H+16311 | HV+16311 | HV+16311 | HV+16311 | HV-16311 | HV-16311 | HV+16311 | B | HV | HV | HV | HV+16311 |
| **CAN22** | H2 | H2 | H | H2 | H2a2a1 | H2a2a1 | H2a2a1 | H30a | H2a2a1 | H27e | H2a2a1 | H | H2a2a1 | H2a2a1 |
| **CAN23** | K1a1b1 | K1a1b1 | K1a1b1 | K1a1b1 | K1a1b1 | K1a1b1 | K1a1b1 | k1a1b1 | K1a1b1 | K1a1b1 | K1a1b1 | K1a1b1 | K1a1b1 | K1a1b1 |
| **CAN24** | K1a14 | K1a14 | K1a14 | K1a14 | K1a14 | K1a14 | K1a14 | K1a12 | K1a14 | K1a14 | K1a14 | K1a14 | K1a14 | K1a14 |
| **CAN25** | J1c2c1 | J1c2c1 | J1c2c1 | J1c2c1 | J1c2c1 | J1c2c1 | J1c2c1 | J1c2c1 | J1c2c1 | J1c2c1 | J1c2c1 | J1c2c1 | J1c2c1 | J1c2c1 |
| **CAN26** | J1c2 | J1c2 | J1c2 | J1c2 | J1c2 | J1c2 | J1c2 | J1c2 | J1c2 | J1c2 | J1c2 | J1c2 | J1c2 | J1c2 |
| **CAN27** | T2c1d+152 | T2c1d+152 | T2c1d+152 | T2c1d+152 | T2c1d+152 | T2c1d+152 | T2c1d-152 | T2c1d-152 | T2c1d+152 | T2c1d2 | T2c1d2 | T2c1d2 | T2c1d | T2c1d+152 |
| **CAN28** | H1cf | H1cf | H1cf | H1cf | H1cf | H1cf | H1 | H1 | H1cf | H1cf | H1cf | H1cf | H1 | H1cf |
| **CAN29** | H6a1a | H6a1a | H6a1a | H6a1a | H6a1a | H6a1a | H6a1a | H6a1a | H6a1a | H6a1a | H6a1a | H6a1a | H6a1a | H6a1a |
| **CAN30** | U5b2b3 | U5b2b3 | U5b2b3a | U5b2b3 | H2a2a1 | H2a2a1 | H2a2a1 | U5b2b3a | H2a2a1 | U5b2b3a | H2a2a1 | U5b2b3a | H2a2a1 | H2a2a1 |
| **CAN31** | U6b1a1 | U6b1a1 | U6b1a1 | U6b1a1 | U6b1a1 | U6b1a1 | U6b1a1 | U6b1a1 | U6b1a1 | U6b1a1 | U6b1a1 | U6b1a1 | U6b1a1 | U6b1a1 |
| **CAN32** | U6b1a | U6b1a | U6b1a | U6b1a | U6b1a | U6b1a | U6b1a | U6b1a | U6b1a | U6b1a | U6b1a | U6b1a | U6b1a | U6b1a |
| **CAN33** | X | X | X3a | X | X | X | X | X3a | X | X3a | X | X3a | X | X |
| **CAN34** | L3f1b1a | L3f1b1a | L3f1b1a | L3f1b1a | L3f1b1a | L3f1b1a | L3f1b1a | L3f1b1a | L3f1b1a | L3f1b1a | L3f1b1a | L3f1b1a | L3f1b1a | L3f1b1a |
| **CAN35** | L3b1a+@16124 | L3b1a+@16124 | L3b1a+@16124 | L3b1a+@16124 | L3b1a+@16124 | L3b1a+@16124 | L3b1a-@16124 | L3b1a-@16124 | L3b1a+@16124 | L3b1a11 | L3b1a | L3b1a | L3b1a | L3b1a+@16124 |
| **CAN36** | U6b1a1 | U6b1a1 | U6b1a1 | U6b1a1 | U6b1a1 | U6b1a1 | U6b1a1 | U6b1a1 | U6b1a1 | U6b1a1 | U6b1a1 | U6b1a1 | U6b1a1 | U6b1a1 |

**Table S5.** BGLMM model results for WGS and WES datasets. Summary of values obtained with the haplogroup classification tool as fixed factor. Mapped reads, mapping quality, and base quality were introduced as covariates. Discordance was used as the response variable and the sample was used as a random factor. The statistical significance is marked with asterisks: * $p \leq 0.05$ | ** $p \leq 0.01$ | *** $p \leq 0.001$

| *Application* | *Tool* | *Estimate* | *Std.Error* | *zvalue* | *Pr(>\|z\|)* | |
|---|---|---|---|---|---|---|
| *WGS* | Haplotracker FASTA | -3.16 | 0.66 | -4.81 | 0.00 | *** |
| - | Phy-Mer FASTA | 0.31 | 0.59 | 0.52 | 0.60 | |
| - | EMMA FASTA | -2.20 | 2.41 | -0.91 | 0.36 | |
| - | Haplofind FASTA | 0.56 | 0.57 | 0.98 | 0.33 | |
| - | MitoTool FASTA | 1.13 | 0.55 | 2.05 | 0.04 | * |
| - | Haplogrep FASTA | -2.18 | 2.35 | -0.93 | 0.35 | |
| - | JamesLick's FASTA | -2.16 | 2.32 | -0.93 | 0.35 | |
| - | HaploCheck BAM | -2.20 | 2.39 | -0.92 | 0.36 | |
| - | MixEmt BAM | 3.55 | 0.66 | 5.35 | 0.00 | *** |
| - | MitoSuite BAM | 0.76 | 0.56 | 1.36 | 0.17 | |
| - | Phy.mer BAM | 0.55 | 0.57 | 0.96 | 0.34 | |
| - | HaploCheck VCF | -2.14 | 2.27 | -0.94 | 0.35 | |
| - | Haplogrep VCF | -2.13 | 2.25 | -0.95 | 0.34 | |
| - | HaploGrouper VCF | -0.43 | 0.71 | -0.61 | 0.54 | |
| - | Mapped Reads | 0.37 | 0.30 | 1.23 | 0.22 | |
| - | Mapping Quality | 0.06 | 0.32 | -1.86 | 0.06 | . |
| - | Base Quality | -0.09 | 0.51 | 0.17 | 0.87 | |
| *WES* | Haplotracker FASTA | -6.84 | 2.82 | -2.42 | 0.02 | * |
| - | Phy-Mer FASTA | 0.47 | 0.48 | 0.98 | 0.33 | |
| - | EMMA FASTA | -0.36 | 0.51 | -0.70 | 0.48 | |
| - | Haplofind FASTA | 0.22 | 0.49 | 0.45 | 0.66 | |
| - | MitoTool FASTA | 0.97 | 0.49 | 1.98 | 0.05 | * |
| - | Haplogrep FASTA | -0.55 | 0.52 | -1.07 | 0.29 | |
| - | JamesLick's FASTA | 0.14 | 0.49 | 0.28 | 0.78 | |
| - | HaploCheck BAM | -4.38 | 1.99 | -2.20 | 0.03 | * |
| - | MixEmt BAM | -1.13 | 0.50 | -2.25 | 0.02 | * |
| - | MitoSuite BAM | -0.63 | 0.48 | -1.32 | 0.19 | |
| - | Phy.mer BAM | 0.27 | 0.45 | 0.60 | 0.55 | |
| - | HaploCheck VCF | -0.59 | 0.52 | -1.13 | 0.26 | |
| - | Haplogrep VCF | -0.57 | 0.52 | -1.09 | 0.27 | |
| - | HaploGoruper VCF | -0.13 | 0.50 | -0.26 | 0.80 | |
| - | Mapped Reads | -5.96 | 3.55 | -1.68 | 0.09 | . |
| - | Mapping Quality | 1.16 | 0.39 | 2.97 | 0.00 | ** |
| - | Base Quality | 0.15 | 0.24 | 0.61 | 0.54 | |
